# Cortical beta coherence provides a stronger non-invasive predictor of movement vigor than local beta power

**DOI:** 10.1101/2025.09.23.678077

**Authors:** Emeline Pierrieau, Claire Dussard, Axel Plantey-Veux, Cloé Guerrini, Nathalie George, Camille Jeunet-Kelway

## Abstract

**Background:** Movement elicits a robust decrease in motor cortical beta-band (β; 13-30 Hz) power contralateral to the moving limb. On this basis, studies have targeted contralateral motor cortical β power to decode or modulate movement vigor (initiation and execution speed) non-invasively. Yet, reported behavioral effects and decoding accuracy remain modest. Considering that controlling vigor involves distributed brain regions, network-level metrics that capture interactions between cortical regions may track changes in vigor more accurately than local power. We therefore tested whether β cortico-cortical coherence, measured as functional connectivity between contralateral motor cortex and other cortical areas, predicts movement vigor more reliably than β power.

**Methods:** Thirty healthy participants performed right hand opening at two instructed speeds (Fast, Slow), while high-density electroencephalography (EEG) was recorded. EEG data were source-localized, and analyses were conducted at the sensor and source levels. We compared β power and β coherence in their ability to discriminate Fast from Slow condition. Effects were assessed across the whole scalp/cortex and using subject-specific selections of electrodes/parcels optimized for discrimination.

**Results:** Fast trials exhibited shorter movement time (MT) and reaction time (RT) than Slow trials, indicating higher vigor. No electrodes cluster showed any significant β power difference between Fast and Slow conditions. With subject-specific channel selection, β power discriminated vigor above chance in most participants, but the polarity of the β power contrast (Fast < Slow or Fast > Slow) varied across individuals. In contrast, with subject-specific parcel selection, β coherence was consistently reduced during Fast relative to Slow in the majority of participants. Across participants, lower β coherence, but not β power, was significantly associated with larger Slow-Fast vigor difference.

**Conclusions:** β cortico-cortical coherence between contralateral motor cortex and other cortical regions provided a more robust and consistent predictor of movement vigor than contralateral motor cortical β power. β coherence exhibited a sustained reduction during fast movements across trials and participants, supporting its use as target for non-invasive neuromodulation of vigor and as feature for decoding intended movement speed.

## Background

Numerous motor disorders involve alterations in the regulation of movement initiation and execution speed, including Parkinson’s disease [1], Huntington’s disease [2], attention deficit hyperactivity disorder [3], and depression [4]. Movement initiation and execution speeds are respectively reflected in reaction time (RT) and movement time (MT) when movement amplitude is kept constant. When incentivized to move fast, RT and MT often co-vary, such that they have been proposed to represent a single behavioral metric called vigor [5,6]. Non-invasive neurostimulation approaches designed to modulate movement vigor have attracted increasing scientific interest. Nevertheless, the development of reliable interventions requires that the underlying brain activity patterns be firmly established as robust predictors of vigor.

Beta-band (β; 13–30 Hz) activity, measurable with electroencephalography (EEG), has emerged as a central marker of movement. The amplitude of β activity (or β power) consistently decreases at central electrodes, located over motor cortical regions, before and during movement [7]. This movement-related β suppression is more pronounced on the side contralateral to the moving limb and can predict the hand selected for movement [8,9]. Thus, β suppression may reflect the activation of underlying motor cortical areas [10]. In line with this, non-invasive modulation of contralateral motor cortical β power has been shown to impact RT and MT, using transcranial alternating current stimulation (tACS) [11,12] and EEG-based neurofeedback (NF) [13–16]. However, reported effects are modest (e.g., ∼30 ms reduction in RT in [15]) and not consistently replicated, as some studies failed to find any significant influence of β modulation on movement speed [17,18]. Furthermore, several studies reported no significant difference in the amplitude of β suppression across movements performed at very different speeds and forces [19–23].

Animal and functional magnetic resonance imagining (fMRI) studies provided evidence that the regulation of movement speed engages distributed networks, including prefrontal, parietal, temporal, occipital, and cingulate cortices, in addition to frontal motor regions [24–26]. Whole-scalp analyses of β power may therefore provide a more comprehensive view of network-level patterns underlying movement vigor. Although prior work has examined the effects of selecting different frequency sub-bands, classifiers, and electrode sites on decoding accuracy, brain activity metrics have typically been restricted to local power [27,28]. Incorporating connectivity measures with source reconstruction of high-density EEG could enable a more precise characterization of network-level dynamics [29]. Coherence is a fundamental connectivity metric that quantifies synchronization between spatially distinct sources of oscillatory activity [30,31]. Imaginary coherence specifically isolates synchronization with a non-zero phase difference (time lag) between signals, thereby reducing artefacts from volume conduction or signal leakage [32]. Robust correlations have been found between RT and cortico-subcortical β coherence, but not β power [33,34]. However, whether cortico-cortical β coherence measured non-invasively can reliably predict movement vigor remains unclear. Furthermore, the correlation reported between cortico-subcortical β coherence and RT was specific to low-β frequencies [33,34]. “β” is in fact not a unitary rhythm: low- and high-β sub-bands have shown opposite sign of correlation with RT [35], consistent with evidence that they originate from distinct neural sources [36]. Most neurostimulation studies aiming to modulate movement speed have targeted ∼20 Hz activity [11,12,15], which may obscure sub-band specific effects.

Therefore, the aim of this study was to identify brain activity patterns that best predict movement vigor across cortical regions, β bands, and metrics (power vs coherence). To do so, we used a high-density EEG dataset (128 channels) collected on a total of 30 participants while they prepared and executed fast and slow hand movements [37]. We measured β power and imaginary coherence (iCoh) in distinct β bands and clusters of electrodes (in sensor space) or brain regions (in source space). We quantified their respective capacity to discriminate movement vigor (fast vs slow), across and within subjects. We found that β iCoh between the motor cortex contralateral to the moving limb and bilateral fronto-parietal regions predicted movement vigor more robustly than contralateral motor β power.

## Methods

The dataset analyzed in the present study corresponds to the experiment 2 data, collected on a total of 30 participants (15 women, 15 men), reported in [37]. In that previous work, the analyses of EEG data focused on assessing power within a broad β band (13–30 Hz) recorded from a single electrode (D19). In contrast, the present study adopts a more comprehensive approach: β power was examined across the entire scalp, subdivided into distinct β sub-bands, and β functional connectivity between brain regions was quantified using source-level iCoh. This extended analysis reflects different research objectives. Whereas Pierrieau et al. (2025) aimed to investigate the role of β power modulation in motor function [37], the present study leveraged this 128-channel EEG dataset to identify neural signatures capable of robustly predicting movement vigor. Details regarding participants and materials are provided in [37]. The experimental design and analysis pipeline are described in detail below.

### Experimental task

#### 1. Overview

Each participant (n = 30) performed two separate sessions, taking place one to three weeks apart from each other. Each session comprised 190 trials, organized in blocks of Comfortable, Fast, and Slow trials (Figure 1A). Participants began each session with one block of 20 Comfortable trials, during which they were asked to execute four repetitions of hand opening at a comfortable, natural pace. The distribution of MT values during this first block was used to determine MT criteria for subsequent Fast and Slow trials in an individualized manner (see Motor task section below). Then, half of participants (n = 15) performed 35 Fast trials, 35 Slow trials, 15 Comfortable trials, 35 Slow trials, 35 Fast trials, and 15 Comfortable trials, in this order, in separate blocks. The other half of participants started with 35 Slow trials, then 35 Fast trials, 15 Comfortable trials, 35 Fast trials, 35 Slow trials, and 15 Comfortable trials. Thus, the order of presentation of blocks of Fast and Slow trials was counterbalanced across participants, but maintained across both sessions from the same participant.

**Figure 1.**
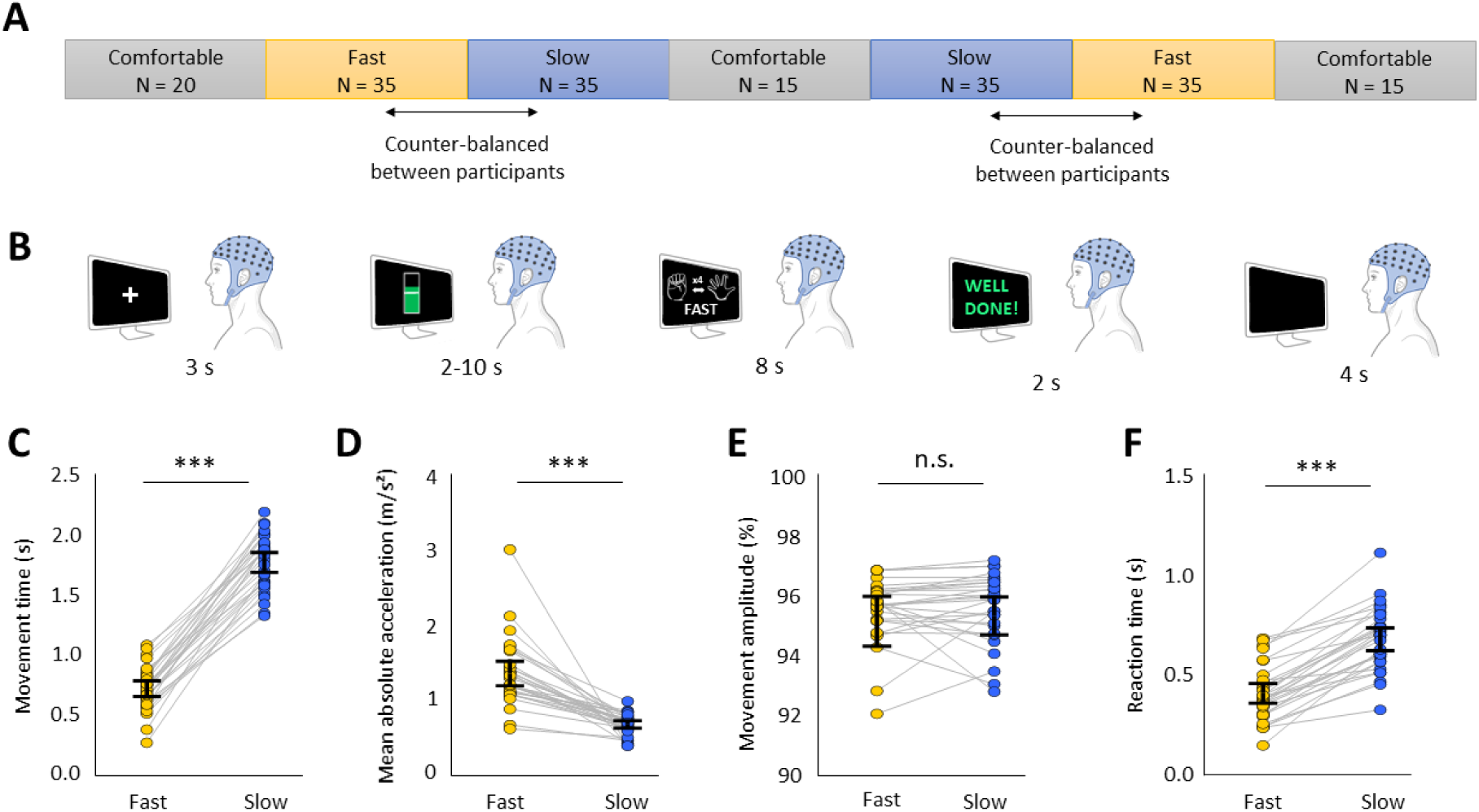
Experimental protocol and behavioral results. **A.** Experiment timeline during each session. The sessions were composed of blocks of trials, here illustrated in a chronological order (from left to right), with speed instruction and number of trials (N) indicated for each block. **B.** Trial timeline. Each trial started with a 3s preparatory period (white fixation cross), followed by a 2 to 10s NF phase with the level of a visual gauge indicating online changes in left motor cortical β power. The motor task started directly after, with the requested speed and number of repetitions of right hand opening recalled to the participants on the screen on each trial. After 8s, the participants received visual feedback about their MT (“Well done!”, “Too fast”, or “Too slow”), such that they could adapt their movement speed on the next trial to reach MT criterion (see Methods). The trial ended with a 4s break with a black screen. **C, D, E, F.** From left to right, average value per participant (dot) and speed instruction (yellow: Fast, blue: Slow) in MT, mean absolute acceleration, movement amplitude, and RT. The vertical black line represents 95% confidence interval around the mean. ***p < 0.001, n.s. = not significant (p > 0.05).

#### 2. Trial timeline

Trial timeline is illustrated on Figure 1B. Each trial began with a white fixation cross presented at the center of the screen for 3 s, instructing participants to remain still and keep their eyes fixated on the cross. In Comfortable trials, the fixation cross was directly followed by the presentation of a go cue indicating the start of the motor task (an illustration of a right hand closed and a right hand open, with an arrow between them and “x4” written above; see Figure 1B). After 8 s, the go cue was replaced by a black screen for 4 s, after which a new trial began with the fixation cross. In Fast and Slow trials, the fixation cross was replaced by a visual gauge reflecting online changes in β power, displayed for 2 to 10 s (see Neurofeedback section). It was immediately followed by the go cue, which was identical to that in Comfortable trials, except that a reminder of the speed instruction (“FAST” or “SLOW”) was displayed below the illustration of the hand. After 8 s, the go cue was replaced by visual feedback about their performance at the motor task for 2 s (see Motor task section). Then, the black screen followed for 4 s, before the beginning of a new trial. Participants were asked to minimize eye and head movements during the whole trial duration, except during the 4 s inter-trial interval (black screen).

#### 3. Motor task

Participants were asked to keep their right hand open, with the back of the hand resting on the table. When the go cue appeared, they were instructed to close their hand four times, performing each movement with maximal amplitude. In Fast and Slow trials, the go cue was followed by visual feedback, which was positive (“Well done!”) if the mean MT of the four repetitions of hand opening reached the speed criterion, and negative if the mean MT fell outside (“Too fast!” / “Too slow”) (Figure 1B). The aim of this feedback was to encourage participants to correct their MT if they started slowing down in Fast trials or speeding up in Slow trials. Speed criteria in Fast and Slow trials were determined based on the distribution of MT values in the first block of trials, which were executed at a comfortable pace. Speed criterion of Fast trials corresponded to mean MT minus 3 standard deviations (SDs), implying that participants received positive feedback whenever their mean MT was equal or inferior to this criterion. In Slow trials, participants received positive feedback whenever their mean MT was equal or superior to mean MT plus 3 SDs.

#### 4. Neurofeedback (NF)

We used a NF paradigm, as detailed in our previous study [26], in order to maximize β power modulations before the motor task. A bidirectional type of NF was designed [39]: participants were trained to regulate motor cortical β power averaged over D19 (C3 equivalent) and adjacent electrodes in opposite directions, either decreasing it (β-down) or increasing it (β-up). These two conditions were carried out separately, in the two sessions of the experiment (one session for β-down, one for β-up). NF was depicted as a virtual gauge, whose level increased as online β power recorded over D19 decreased in β-down and increased in β-up. If participants successfully maintained the gauge at the targeted level for 2 s, NF stopped and was directly replaced by the go cue. NF was automatically replaced by the go cue after 10 s if participants failed to reach this criterion. NF was presented only in the blocks of Fast and Slow trials, not in Comfortable trials. Each block of Fast and Slow trials included 20 trials with real NF, followed by 15 trials with sham NF. This sham NF consisted of a replay of the gauge presented during the 15 first trials of the block, and participants were asked not to try controlling the level of the gauge. This passive control condition served the purpose of our initial study [37] and was not relevant here. Further details on the NF setup can be found in [37].

In the present study, trials from all NF conditions (β-down, β-up, and passive control) were pooled together and not distinguished further in the analyses. We verified that NF duration did not significantly differ between Fast and Slow trials (t(29) = -0.7, p = 0.465, d = - 0.14), such that it could not account for differences in behavior and β activity between Fast and Slow trials.

### Data processing

Behavior (luxmeter and accelerometer) and EEG data processing was conducted on Matlab R2023A (MathWorks) and Python 3.10.11 within Spyder 6.0.3 environment.

#### 1. Behavior

MT, RT and movement amplitude were derived from the signal of a light sensor (LUX, BITalino) positioned at the palmar surface of the third digit fingertip of the right hand. MT was computed as the time separating 2 luminosity peaks detected by the luxmeter, corresponding to the moments when the hand was fully open. MT was averaged across the 4 hand movement repetitions of each trial. Movement amplitude was estimated as the difference between the maximal and minimal luxmeter values per repetition (data points starting from the detected luminosity peak, until the data point preceding the next luminosity peak), averaged across the 4 repetitions of each trial. RT was computed as the time separating go cue onset and the first luminosity peak. Acceleration was recorded using an accelerometer (ACC, BITalino) positioned on the dorsal side of the third digit fingertip of the right hand. Mean absolute acceleration was measured by averaging absolute data from the accelerometer signal between go cue onset and the end of the fourth repetition (fourth luminosity peak). Outliers were calculated separately for each motor variable per participant and speed instruction. They were defined as values inferior to the median minus 3 median absolute deviations (MADs) or superior to the median plus 3 MADs. Trials that were identified as outliers were marked as NaNs and not considered further in the analyses.

#### 2. EEG

##### a. Preprocessing

Preprocessing of raw EEG data was performed using EEGLAB v2024.2.1 in Matlab [40]. First, data was downsampled to 256 Hz. Then, a 1-49 Hz bandpass, two-pass, finite impulse response (FIR) filter was applied. Noisy channels were automatically detected using the Clean raw data algorithm from EEGLAB with the following parameters: signal flat for more than 4 s, standard deviation of high-frequency noise superior to 4, and correlation with nearby channels inferior to 0.85. The original signal from those channels was removed and interpolated based on the activity from their neighboring electrodes. Data was then segmented into epochs of 14 s duration locked around go cue onset (-11.5 s to +2.5 s). This period included the time window of NF presentation and movement preparation and execution, with an additional minimum 1.5 s period before NF presentation, which was used as baseline period for subsequent analyses. Independent component analysis (ICA) was run on all epochs of all conditions, for each participant separately, using runica algorithm from EEGLAB (with pca parameter set to 60). Artefactual components related to eye movements were identified based on their anterior location, spurious occurrences and low frequency dominant spectrum [41] and removed for each participant and session (min number of components removed = 0, max = 2). The signal was then re-referenced to the average of all EEG electrodes, at each time point.

##### b. Computation of β power in the sensor space

Preprocessed EEG data from EEGLAB (.set files) were imported into Spyder environment using MNE Python 1.9.0. Time-frequency decomposition was applied using Morlet wavelets on each epoch, cropped between –1.5 and +1.5s around go cue onset, using .compute_tfr() function (method=”morlet”, freqs=[15:25] with 1Hz step, n_cycles=[15:25]/2, average=False, decim=2, output=”complex”). The resulting data matrix was averaged over the 15 to 25 Hz frequencies. β power was then computed as the squared absolute value of the matrix. Finally, β power was averaged over seven 0.5s time windows centered on go cue onset, with 50% overlap: [-1.00; -0.50], [-0.75; -0.25], [-0.50; 0.00], [-0.25; 0.25], [0.00; 0.50], [0.25; 0.75], [0.50; 1.00]).

The above procedure was applied separately for epochs from Fast and Slow conditions and for each β sub-band (modifying freqs parameter in .compute_tfr() with the following values: low β = 8-15 Hz, standard β = 15-25 Hz, high β = 25-35 Hz). For each condition and β sub-band, we obtained a 140 (trials) x 128 (channels) x 7 (time windows) β power matrix.

The same procedure was applied to compute baseline β power, except that the epochs from Fast and Slow conditions were pooled together and centered on the onset of the visual gauge used for NF and passive control conditions. Baseline β power was averaged over the three 0.5s time windows preceding NF onset ([-1.50; -1.00], [-0.75; -0.25], [-0.50; 0.00], which resulted in a 280 (trials) x 128 (channels) x 3 (time windows) baseline β power matrix.

##### c. Computation of β iCoh and β power in the source space

A separate pipeline was used to compute β power and iCoh in the source space. As for β power, β iCoh was computed on the preprocessed EEG data imported in MNE Python and epoched between -1.7 and +1.7s around go cue onset (the extra 0.2s at the beginning and end of the epochs was added to limit edge artefacts). For each condition (Fast/Slow), epochs were band-pass filtered in the three β sub-bands (low β = 8-15 Hz, standard β = 15-25 Hz, high β = 25-35 Hz) using zero-phase FIR filters and decimated at 128Hz. Filtering was chosen instead of a full time–frequency decomposition because adding an explicit frequency dimension made the data too large to handle with inverse modeling, and averaging across frequencies would have distorted the phase information. Source estimates were obtained per epoch with the apply_inverse_epochs function using a pre-computed inverse operator based on the fsaverage template head model (boundary-element forward model and oct6 source space), a noise covariance matrix computed from −2 to 0 s before NF onset regularized via a shrinkage estimator (method=’shrunk’), minimum norm estimate (MNE) as the inverse method, fixed normal orientations (pick_ori=’normal’), and lambda2=1 (SNR = 1); depth weighting was set to 3. Normal orientation was chosen to limit inter-individual variability that could be induced with free orientations and affect between-subject comparisons of β activity in Fast and Slow conditions. Label-wise time courses were extracted using Desikan-Killiany parcellation [42], and symmetric orthogonalization [43] was applied to reduce zero-lag mixing. Analytic signals were then obtained via the Hilbert transform.

β iCoh was calculated as the magnitude of the imaginary part of the normalized cross-spectrum between the left precentral gyrus, corresponding to the approximate location of the left motor cortex, and each other parcel. β iCoh was then averaged within the same predefined time windows as for β power, spanning −1.0 to +1.0 s around go cue onset, which resulted in 140 (trials) x 67 (parcels) x 7 (time windows) β iCoh matrices for each condition (Fast, Slow) and each β sub-band (low, standard, high β). Similar to sensor-level β power, source-level β power was calculated as the squared absolute value of the time-frequency matrix, before being averaged within the same predefined time windows. This resulted in 140 (trials) x 68 (parcels) x 7 (time windows) β power matrices for each condition (Fast, Slow) and β sub-band (low, standard, high β).

### Data analysis and statistics

Data analysis was conducted using Python 3.10.11 within Spyder 6.0.3 environment. The analyses displayed in the main manuscript are those conducted on the standard β sub-band (15-25 Hz). The analyses on the other β sub-bands (low and high β) are presented as Supplementary Results.

#### 1. Motor variables

Motor variables (MT, RT, movement amplitude, mean absolute acceleration) were averaged across trials per subject and compared between Fast and Slow conditions. When data distribution followed a normal distribution (Shapiro test p-value > 0.05), statistical comparisons were performed using paired t-tests, otherwise Wilcoxon rank tests were applied. Effect sizes were reported as Cohen’s dz (d) for paired t-tests results, and as rank biserial correlation (r) for Wilcoxon rank tests.

#### 2. Between-subject, whole scalp/brain analyses of β power and β iCoh

For between-subject analysis, data was averaged across trials. For each subject and β sub-band, trials with extreme β power values were first identified using a MAD-based outlier rejection (|x−median| > 3×MAD) per channel and time window and removed before averaging across trials (6.2% trials removed in average). Baseline values were processed the same way and then averaged across the three pre-NF time windows and replicated over all time windows to serve as a time-invariant comparator.

Cluster-based permutation tests were run at the sensor level to compare Fast and Slow conditions to each other and against baseline, in each β sub-band and time window. For this, we concatenated the β sub-band power values obtained on every electrode in each time window across subjects, resulting in paired matrices (30 subjects x 128 channels) for the Fast and Slow conditions in each time window. Baseline data were similarly concatenated across subjects. A sensor adjacency graph was built from 2D EEG positions, connecting electrodes within 0.02 Euclidean units. It was then used in mne.stats.permutation_cluster_test() function with two-sided paired t-testing, a critical threshold set for alpha = 0.05, 1000 permutations, and the sensor adjacency graph to define clusters. To facilitate their interpretation, β power in Fast and Slow conditions was then baseline-corrected per subject, to convert it into dB (10*log(β power/baseline β power)). Mean difference in baseline-corrected β power between Fast and Slow across subjects and 95% confidence intervals were computed and averaged over channels of interest.

With regard to β iCoh, for each subject and β sub-band, trials with extreme β iCoh values were first identified using a MAD-based outlier rejection (|x−median| > 3×MAD) per parcel and time window and removed before averaging across trials (2.3% trials removed in average). Paired t-tests were computed to compare β iCoh between Fast and Slow conditions in each parcel and time window. Considering the high number of comparisons (67 parcels * 7 time windows = 469 tests), p-values were not interpreted, and topographical maps of t-values were used as a qualitative tool to assess the patterns of β iCoh modulations across the brain. In addition, the five parcels with the highest absolute t-values, averaged across all time windows, were represented as a bar plot in order to visualize their location and the sign of modulation in the brain areas where the functional connectivity (β iCoh) with the left precentral gyrus was the most different in Fast compared to Slow conditions. Mean difference in β iCoh between Fast and Slow conditions across subjects and 95% confidence intervals were computed and visualized for these top five regions of interest.

#### 3. Subject-specific channel/parcel selection using logistic regressions

Channel/parcel selection for analyzing β activity can be conducted subject-wise, focusing on channels/parcels that provide the strongest predictive signals [38,39]. This method accounts for inter-individual differences in the spatial distribution of speed-related changes in β activity. For each subject, we used logistic regression with L1 regularization, embedded in a 5-fold stratified cross-validation scheme. β activity (β power or β iCoh) was used as predictor, and speed instruction (Fast/Slow) as the outcome to predict. Within each training fold, trials were first cleaned separately per condition using a MAD-based outlier rejection (|x−median| > 3×MAD), and features were standardized (z-scored) before model fitting. Feature selection was performed using sequential forward selection (SFS): starting from no features, we added spatial groups of features one by one if they improved the mean cross-validated AUC by at least 0.002. In sensor-space analyses, each group of features corresponded to a single EEG channel, whereas in source-space analyses, it corresponded to a single parcel; in both cases, all associated time windows were included. Including all time windows not only simplified the interpretation of features selection but also enabled the identification of temporally stable features, that could discriminate Fast from Slow trials consistently across time. Such features could serve as reliable targets for neurostimulation applications, considering that they most often modulate brain activity over several seconds (e.g., NF and tACS). L1 regularization further allowed different time windows within each features group to be weighted according to their predictive value. While L1 regularization acted within features groups, the SFS procedure constrained the number of features groups at a higher level. To prevent overly complex models and improve interpretability, a maximum of three groups of features was retained per participant, corresponding to 21 features in total (3 channels/parcels * 7 time windows). Once the optimal subset of features was identified, we assessed its stability and generalization by bootstrap resampling. For each participant, we created 1,000 bootstrap samples by randomly drawing trials from their dataset with replacement. On each bootstrap sample, we ran 5-fold cross-validation and computed the mean AUC. From the distribution of AUC values across the 1,000 bootstrap samples, we then derived model performance and its 95% confidence interval.

Then, we examined the consistency of the selected features at the group level. Participants whose confidence interval overlapped chance level (AUC = 0.5) were excluded from group-level aggregation. For the remaining participants, the most selected features were represented as topographical heatmaps, in order to assess whether some specific channels or parcels consistently predicted speed instruction across participants. In addition, their standardized regression coefficients were averaged across cross-validation folds and selected features, with the aim to determine if the dominant direction of modulation of the selected features with movement speed (i.e., higher or lower in Fast vs Slow) was consistent across participants. To do so, a one-sample t-test against zero was applied to the mean regression coefficients, obtained after averaging regression coefficients across all selected features per participant. The percentages of participants showing positive and negative mean regression coefficients were also extracted.

#### 4. Between-subject correlation analyses of regression coefficients with movement vigor

We ran correlation tests across subjects between mean regression coefficients from within-subject models (see Section 3 above) and Slow – Fast difference in vigor, computed as the sum of RT and MT. Correlations were run separately for mean regression coefficients based on β power and those based on β iCoh data. Pearson’s r correlation coefficients are reported in the text, along with p-values from one-sided tests. One-sided t-tests were used since a negative correlation was expected between β power and vigor, considering that previous evidence linked faster movement initiation and execution to lower β power [11,15,40]. A similar relationship has been reported between cortico-subcortical coherence and RT [33,34]. We further ran correlation tests of Slow – Fast difference in MT and RT separately, with mean regression coefficients based on β power and those based on β iCoh, in order to determine if correlations with vigor were mostly driven by differences in MT or RT.

## Results

### 1. Impact of speed instruction (Fast/Slow) on movement vigor

Mean MT was strongly affected by instruction speed across participants, as demonstrated by significantly shorter MT in Fast in comparison to Slow conditions (t(29) = -24.3, p < 10^-16^, d = -4.43) (Figure 1C). This effect was not explained by a significant change in movement amplitude between Fast and Slow conditions (W(29) = 168, p = 0.191, r = -0.28) (Figure 1D), and mean absolute acceleration was significantly higher in Fast than in Slow conditions (W(29) = 465, p = 10^-8^, r = 1.00) (Figure 1E). RT was significantly shorter in Fast than in Slow conditions (t(29) = -13.3, p = 10^-13^, d = -2.42) (Figure 1F). Together, these results confirmed that speed instruction significantly impacted movement vigor, Fast condition being associated with higher movement vigor than Slow condition.

### 2. β power modulation according to speed instruction

β power was first computed in a ±5 Hz range around 20 Hz (15-25 Hz), considering that these frequencies have been the most targeted by neurostimulation studies aiming at modulating movement speed [11,12,15]. β power was measured from -1 s to +1 s around the go cue onset to study both proactive (before go cue) and reactive (after go cue) adjustments. Indeed, under the block design, participants could prepare in advance of the go cue, knowing whether the forthcoming movement would be Fast or Slow (Figure 1A). β power was averaged in 0.5 s windows with 50% (0.25 s) overlap, similar to the method used in previous studies [15,41]. β power was first analyzed at the group level across the whole scalp, using cluster-based permutation tests.

When compared to baseline, β power was decreased after go cue onset both in Fast and Slow conditions, with significant clusters observed between 0.25 and 1s, starting over bilateral occipital electrodes and extending to parietal and central electrodes, especially over the left hemisphere (Fast vs baseline: tmean = [-3.52; -2.78], p = [0.015; 0.026]; Slow vs baseline: tmean = [-3.53; -2.58], p = [0.016; 0.044]) (Figure 2A). This finding confirmed that fast and slow movements both produced a significant β event-related desynchronization (ERD), corresponding to the pattern often targeted in BCI and neurostimulation studies aiming to decode or modulate movement speed [13,15,21,28]. Yet, cluster-based permutation tests did not reveal any significant difference in β power across all channels when comparing Fast and Slow conditions, suggesting that the magnitude of the movement-related β suppression was not significantly impacted by speed instruction. Likewise, low β (8-15 Hz) power and high β (25-35 Hz) power significantly decreased following go cue onset compared to baseline, but they did not significantly differ between Fast and Slow conditions (Figures S1A, S2A).

**Figure 2.**
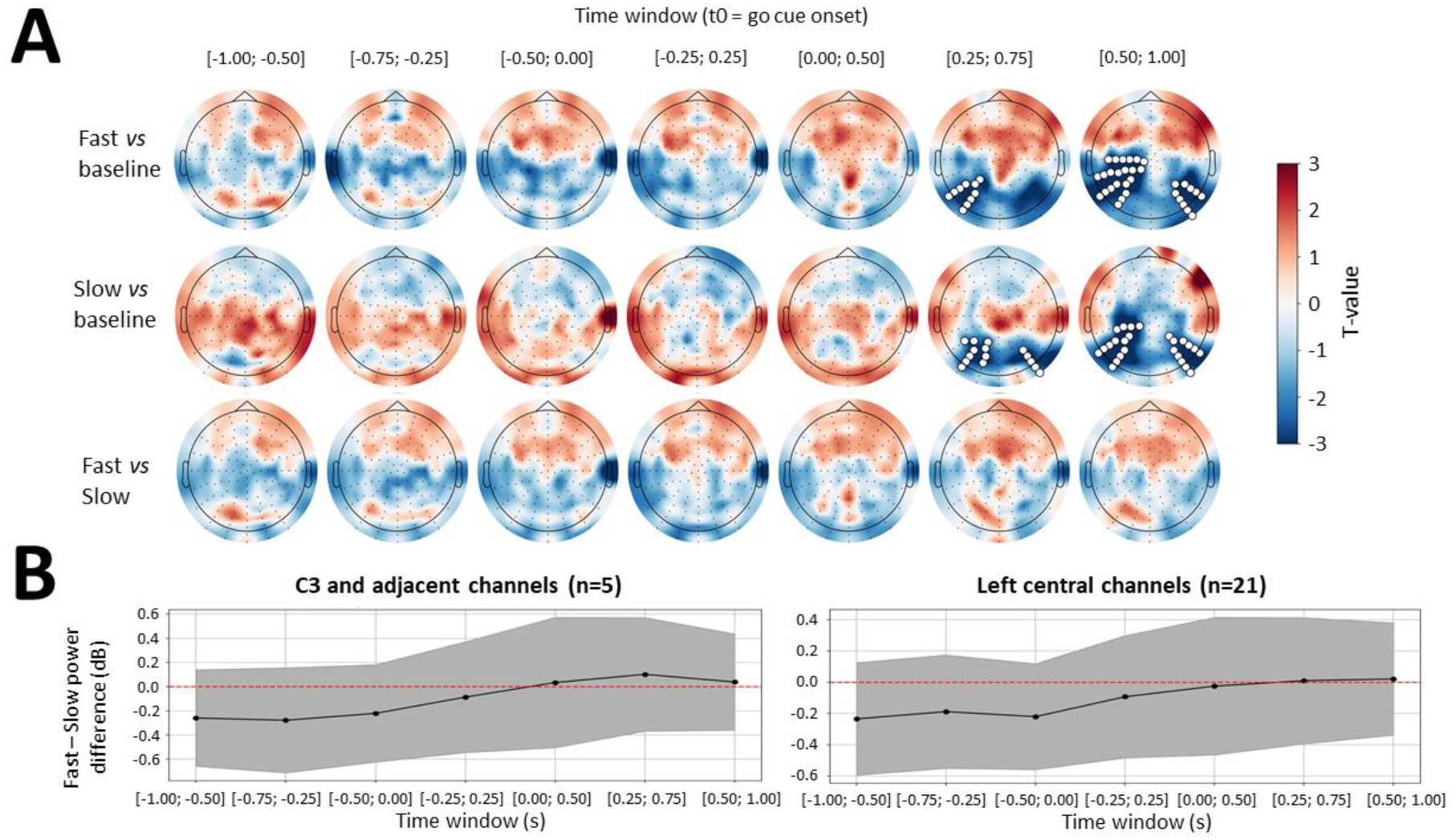
Analysis of mean difference in β power between Fast and Slow conditions across the whole scalp and over left central electrodes. **A.** Topographical maps of t-values resulting from cluster-based permutation tests, which compared β power in Fast to baseline (top row), β power in Slow to baseline (middle row), and β power in Fast to Slow (bottom row). The mean difference in β power was assessed in each 0.5s time window separately, that are indicated on top of each plot, in time relative to go cue. Electrodes that belonged to significant clusters (p < 0.05) are in highlighted in white. **B.** Mean (dot) and 95% confidence intervals (shaded area) of the difference between Fast and Slow conditions in β power, depicted per time window (x-axis), averaged over C3 and adjacent electrodes (D19, D12, D28, D18, D20; left) and over a larger subset of left central electrodes (D1, D15, D16, D2, D14, D13, D18, D17, D19, D20, D12, D18, D28, D27, D11, D10, D21, D26, D29, A6, A7; right). (n= …) indicates the number of electrodes included in each subset.

We further analyzed whether speed instruction produced localized changes in β power at left central electrodes. Analyses were conducted on β power over a small subset of left central electrodes (D19, D12, D28, D18, D20), corresponding to C3 and adjacent electrodes, which are often targeted in protocols aiming to modulate movement initiation and execution speed [13,15,16], and over a larger subset of left cluster electrodes (D1, D15, D16, D2, D14, D13, D18, D17, D19, D20, D12, D18, D28, D27, D11, D10, D21, D26, D29, A6, A7), closer to the spatially broad reduction in β power found when comparing Fast and Slow conditions to baseline (Figure 2A). Although mean β power was overall lower in Fast than Slow conditions during the pre-go cue period in both groups of electrodes, the difference in β power between Fast and Slow conditions did not reach statistical significance in any time window for either group (Figure 2B). Similar results were found for low and high β power (Figures S1B, S2B).

For β power to serve as an effective target for neurostimulation and decoder of movement speed, it should reliably discriminate between Fast and Slow conditions both at the group level (across subjects) and at the individual level (across trials). Additionally, group averages may mask subject-specific effects, particularly in relation to electrode preselection. The scalp topography of movement-related changes in β power can vary across individuals, which is why many BCI studies perform subject-specific electrode selection based on the electrodes that best predict movement [38,39]. Following this approach, we computed, for each participant, a logistic regression with speed instruction (0 = Fast, 1 = Slow) as the outcome and single trial β power as predictor. β power was first extracted from the large left central subset of electrodes previously identified (D1, D15, D16, D2, D14, D13, D18, D17, D19, D20, D12, D18, D28, D27, D11, D10, D21, D26, D29, A6, A7), in all time windows. We first applied sequential forward selection (SFS) to constrain the number of features included in the model (maximal number of channels selected = 3) (see Methods for details). The accuracy of model predictions was determined by computing bootstrapped (n = 1000 permutations) 95% confidence intervals of cross-validated area under the curve (AUC) scores.

AUC results showed above-chance performance (i.e., lower bound of 95% CI > 0.5) for 26 of 30 participants (Figure 3A). Thus, models identified β power patterns that could distinguish Fast from Slow conditions in most subjects. The distribution of the electrodes selected in the models that showed above-chance performance was skewed toward the superior and lateral regions of the left central cluster, with D19 (C3 equivalent) not being often selected among participants (Figure 3B). We then computed the mean regression coefficient across selected features per participant and assessed whether it significantly differed from zero, which would indicate consistent direction of left motor cortical β power modulation across subjects. The mean regression coefficient did not differ significantly from zero (t(25) = 0.43, p = 0.673, d = 0.08), with 14/26 (53.8%) participants showing negative and 12/26 (46.2%) positive values (Figure 3C). This near-even split indicates low inter-individual consistency in the direction of left motor cortical β power modulation according to speed instruction (Fast vs Slow).

**Figure 3.**
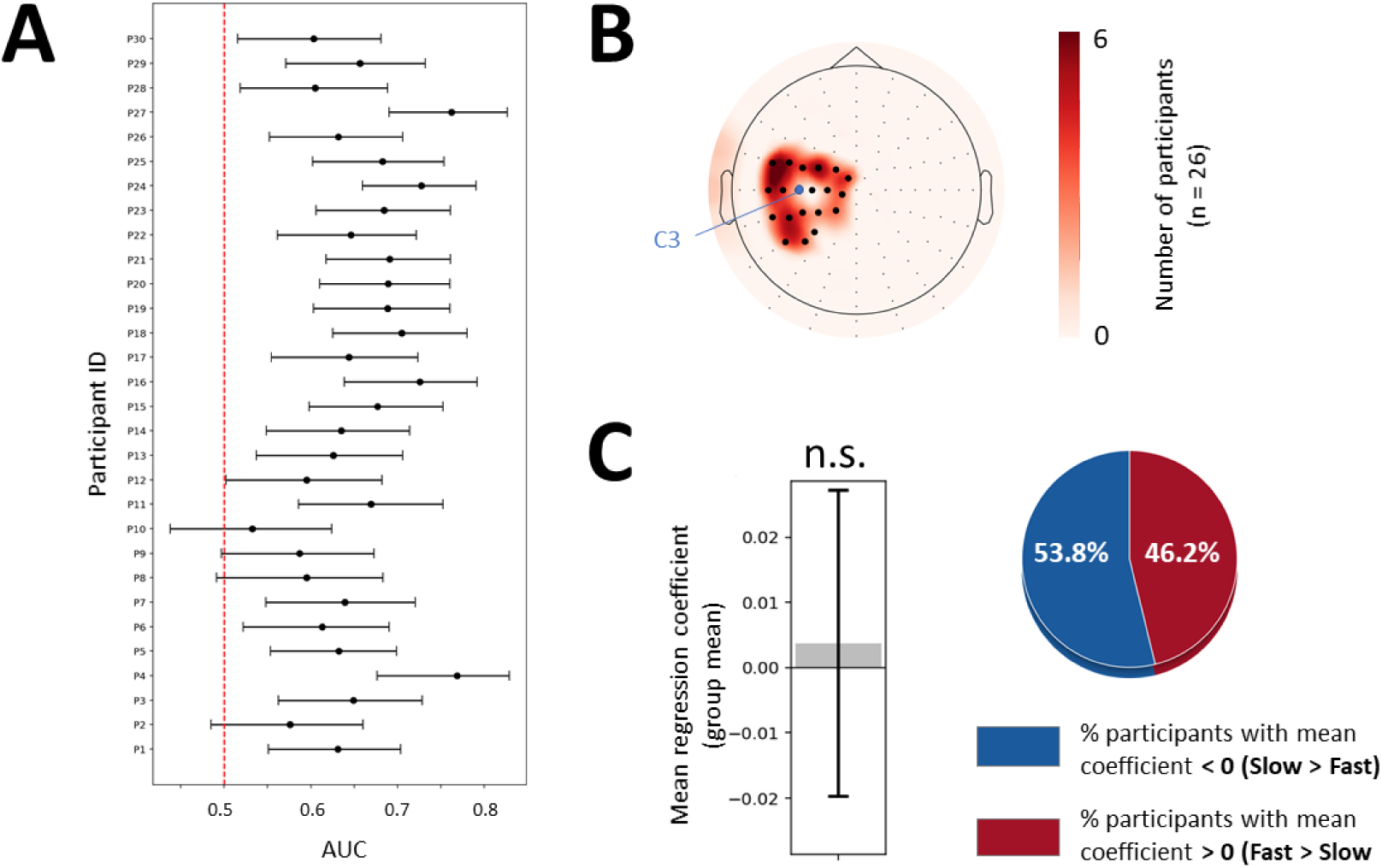
Analysis of mean difference in β power between Fast and Slow conditions with subject-specific selection of left central electrodes. **A.** 95% confidence intervals around the mean AUC represented as error bars for each participant (y-axis) resulting from logistic regression models of speed instruction (Fast vs Slow) based on left central β power. The red dotted line indicates chance level (AUC = 0.5). **B.** Scalp topography of left central electrodes according to their selection frequency. Red areas indicate electrodes most often selected across participants. C3 (D19) is marked with a blue dot. Electrodes included as candidate features are highlighted with thick black dots. **C.** Left, mean regression coefficient (averaged across subjects) represented as a gray bar, with the error bar illustrating 95% confidence interval. N.s. = not significant (p > 0.05). Right, pie chart illustrating the percentage of participants showing positive and negative regression coefficients in red and blue, respectively.

Restricting the features included in the model to pre-movement time windows ([-1.00; -0.50], [-0.75; -0.25], [-0.50; 0.00], [-0.25; 0.25]) – considering that the group difference in β power between Fast and Slow conditions was in average more pronounced during this period (Figure 2B) – did not significantly improve the global performance of the models, with 25 out of 30 participants showing above-chance performance, and mean regression coefficient still not significantly different from zero (t(24) = -0.67, p = 0.508, d = -0.13) (Figure S3A-C). Alternatively, keeping all time windows and extending β power features to all electrodes (excluding the most peripheral ones to limit the influence of residual artefacts) led to better model accuracy, but the mean regression coefficient remained not significantly different from zero (t(29) = -1.35, p = 0.188, d = -0.25) and selected electrodes were mostly located in the periphery (Figure S3D-F). Finally, applying the same modeling procedure as for the initial analysis on standard β power (i.e., left central electrodes, all time windows) to low and high β power also did not improve model accuracy and resulted in variable sign of left motor central β power modulation across subjects (low β: t(19) = -0.35, p = 0.732, d = -0.08; high β: t(27) = 0.19, p = 0.854) (Figure S4). Hence, no consistent sign of β power modulation with movement speed was identified across participants, regardless of electrode grouping or β sub-band selection.

### 3. β iCoh modulation according to speed instruction

EEG data was source reconstructed and averaged in parcels following the Desikan-Killiany atlas (see Methods for details). Then, β iCoh between the left precentral (motor) cortex and all the other parcels was computed to estimate β activity synchrony between these regions. β iCoh was averaged within the same time windows and β bands as the ones used in previous β power analyses. β iCoh appeared overall lower in bilateral fronto-parietal regions when comparing Fast to Slow conditions (Figure 4A). This tendency was confirmed by the observation that the parcels with the highest absolute t-values (averaged across time windows) were all located in frontal and parietal regions, and most often (4/5 parcels) showed negative t-values, indicating decreased β iCoh in Fast as compared to Slow conditions (Figure 4B). Although not significantly different from zero, all these parcels consistently showed lower β iCoh in Fast than Slow conditions across time windows, both before and after go cue with a momentary increase at go cue onset, except for β iCoh between left precentral and left postcentral parcels, which tended to increase after go cue (Figure 4C). Similar analyses conducted on low and high β iCoh displayed inconsistent direction of modulation when comparing β iCoh in Fast to Slow conditions across time windows (Figure S5).

**Figure 4.**
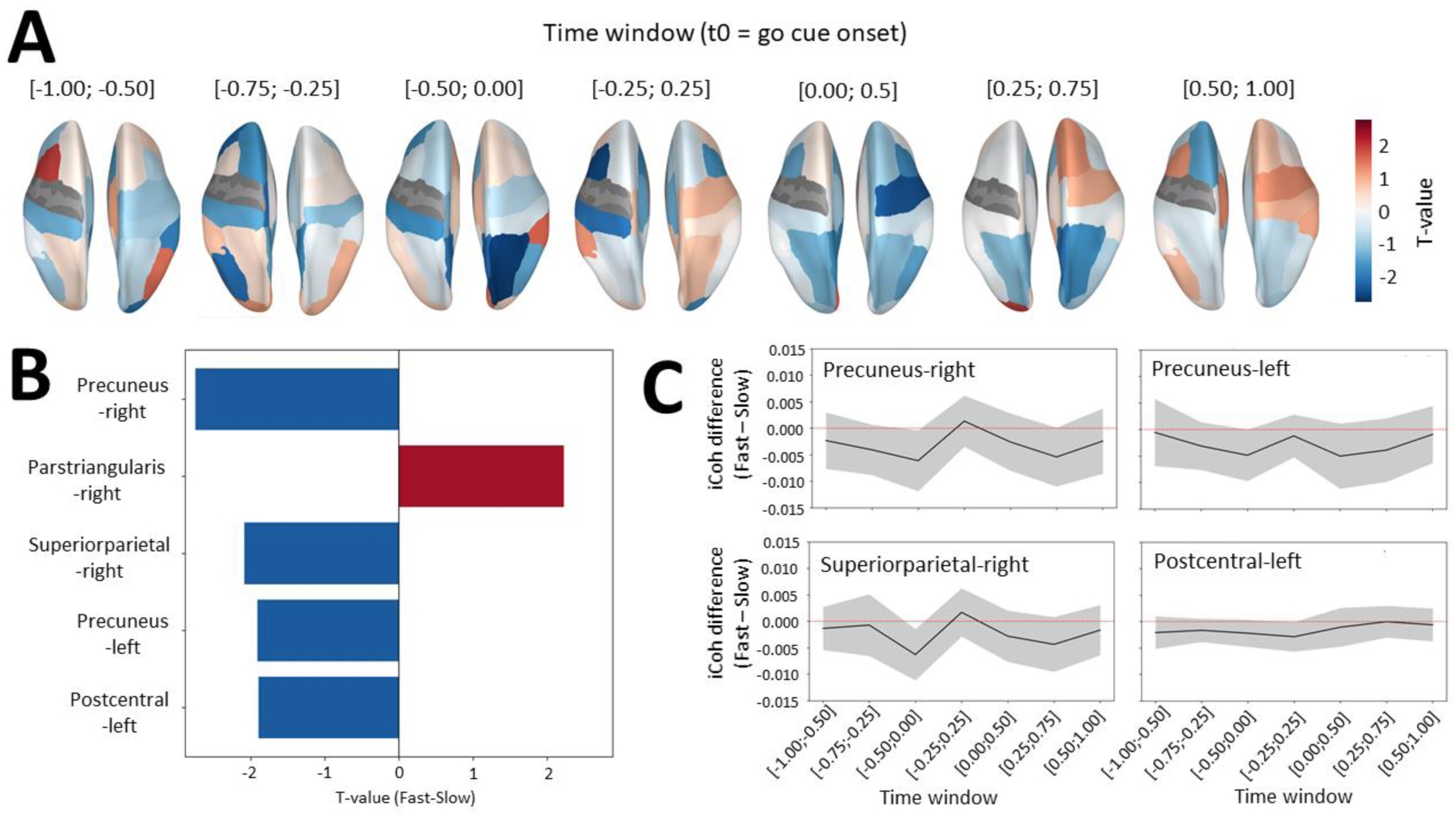
Analysis of mean difference in β iCoh between Fast and Slow conditions across parcels. **A.** Topographical maps of t-values resulting from the comparison of β iCoh in Fast and Slow conditions. Mean difference in β power was assessed in each 0.5s time window separately, that are indicated on top of each plot, in time relative to go cue. Hot and cold colors respectively indicate higher (positive t-value) and lower (negative t-value) β iCoh in Fast than Slow conditions in average. **B.** Top five parcels with the highest absolute t-value averaged across time windows. Red and blue bars respectively indicate parcels associated with a positive and negative t-value. **C.** Mean and 95% confidence intervals (shaded area) of the Fast minus Slow difference in β iCoh, depicted per time window, for the four parcels with the most negative t-value illustrated in Figure 4B (precuneus-right, precuneus-left, superiorparietal-right, postcentral-left).

β iCoh difference between Fast and Slow conditions was also assessed using subject-specific parcel selection, applying the same procedure as for β power at the source level. For each participant, we computed a logistic regression across trials, with speed instruction associated with each trial (0 = Fast, 1 = Slow) as the outcome and trial-by-trial β iCoh between the left motor parcel and bilateral fronto-parietal parcels as predictors. 21 bilateral (left and right) fronto-parietal parcels were included as predictors in the model, in each time window: precuneus, superior parietal, inferior parietal, supramarginal, postcentral, precentral (right only), superior frontal, caudal middle frontal, rostral middle frontal, pars opercularis, pars triangularis. Note that the number of parcels included (n = 21) matches the number of channels included in the analysis of sensor-level β power. The same settings were also applied for SFS (maximal number of parcels selected = 3) and AUC estimation.

AUC results showed above-chance performance for 28 out of 30 participants (Figure 5A). Thus, similar to β power, models identified β iCoh patterns that could discriminate Fast from Slow conditions in most subjects. parcels selected in the models that showed above-chance performance (n = 28) were distributed across bilateral fronto-parietal regions, with right precuneus and postcentral parcels being the most often selected (Figure 5B). Mean regression coefficient was significantly lower than zero (t(27) = -2.32, p = 0.028, d = -0.44), indicating that the selected parcels most often showed lower β iCoh in Fast than Slow conditions, with 20/28 (71.4%) participants showing negative and 8/28 (28.6%) positive values (Figure 5C).

**Figure 5.**
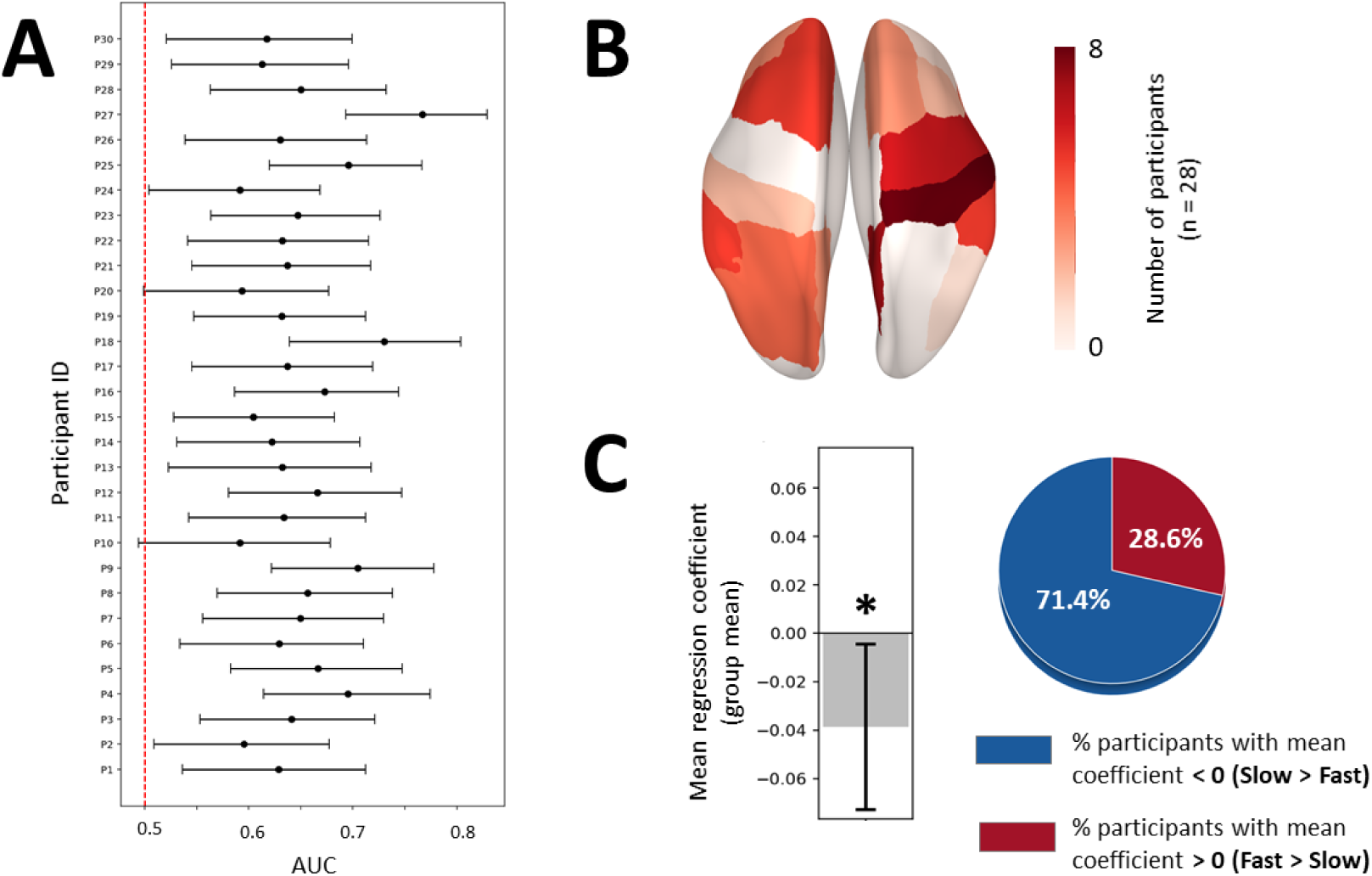
Analysis of mean difference between Fast and Slow conditions in left precentral β iCoh with subject-specific selection of bilateral fronto-parietal parcels. **A.** 95% confidence intervals around the mean AUC represented as error bars for each participant (y-axis) resulting from logistic regression models of speed instruction (Fast vs Slow) based on fronto-parietal β iCoh. The red dotted line indicates chance level (AUC = 0.5). **B.** Brain topography of fronto-parietal parcels according to their selection frequency. Red areas indicate parcels most often selected across participants. **C.** Left, mean regression coefficient (averaged across subjects) represented as a gray bar, with the error bar illustrating 95% confidence interval. *p < 0.05. Right, pie chart illustrating the percentage of participants showing positive and negative regression coefficients in red and blue, respectively.

The same analysis pipeline was applied to low and high β iCoh. Mean AUC score across participants was not significantly different using low β iCoh (t(29) = -0.73, p = 0.470, d = -0.13) or high β iCoh (t(29) = -1.14, p = 0.263, d = 0.21) as predictor in comparison to using standard β iCoh. As for standard β iCoh, the vast majority of participants showed above-chance model performance (Figure S6A, D). Mean regression coefficient was not significantly different from zero for the two bands (low β: t(27) = 0.32, p = 0.750, d = 0.06; high β: t(24) = -0.74, p = 0.468, d = -0.15), with near-even split of positive and negative coefficients across participants (Figure S6C, F).

Differences in the stability of regression coefficients between models based on β power and those based on β iCoh may partly reflect that the two metrics were estimated in different spaces. β power was originally computed in sensor space, whereas β iCoh was derived from source-reconstructed signals. Source reconstruction acts as a spatial filter and can improve statistical detectability by reducing signal mixing caused by volume conduction [42]. Consequently, sensor-level β power models may be more prone to capturing noise and yield less stable coefficients across participants. To test this, we repeated the full pipeline at the source level for β power. Similar to β iCoh, the vast majority of participants showed above-chance model performance (Figure S7A). Parcels selected in the models that showed above-chance performance (n = 28) were distributed across bilateral fronto-parietal regions (Figure S7B). However, consistent with sensor-level β power results, mean regression coefficient did not differ from zero (t = -0.13, p = 0.895, d = -0.03), with an approximately even split of positive and negative coefficients across participants (Figure S7C).

### 4. Correlation analysis between mean regression coefficients and movement vigor

Between-subject correlation analyses were conducted to test whether mean regression coefficients were associated with inter-individual Slow – Fast differences in movement vigor, computed as the sum of RT and MT. Given the within-participant predominance of negative β-band iCoh coefficients for Fast versus Slow, we assessed whether this relation generalizes across participants, that is, whether more negative β iCoh coefficients predict larger Slow – Fast vigor differences. Analogous analyses were conducted for β power. Based on prior reports linking faster movement initiation and execution to lower contralateral motor β power [11,15,40] and on the present observation of reduced β iCoh in Fast compared to Slow conditions, we predicted negative associations for both metrics.

Consistent with this prediction, we found a significant negative correlation between inter-individual vigor differences (Slow – Fast) and mean regression coefficients based on β iCoh (r = –0.33, p = 0.038) (Figure 6A). In contrast, mean regression coefficients based on β power were not significantly correlated with inter-individual vigor differences (r = –0.11, p = 0.286) (Figure 6C). Specifically, a significant negative association was found between mean regression coefficients based on β iCoh and inter-individual differences in MT (r = –0.38, p = 0.020) (Figure 6B), but not RT (r = –0.16, p = 0.197) (Figure 6C). No significant association was observed between mean regression coefficients based on β power and inter-individual differences in MT (r = –0.13, p = 0.246), but not RT (r = –0.04, p = 0.422).

**Figure 6.**
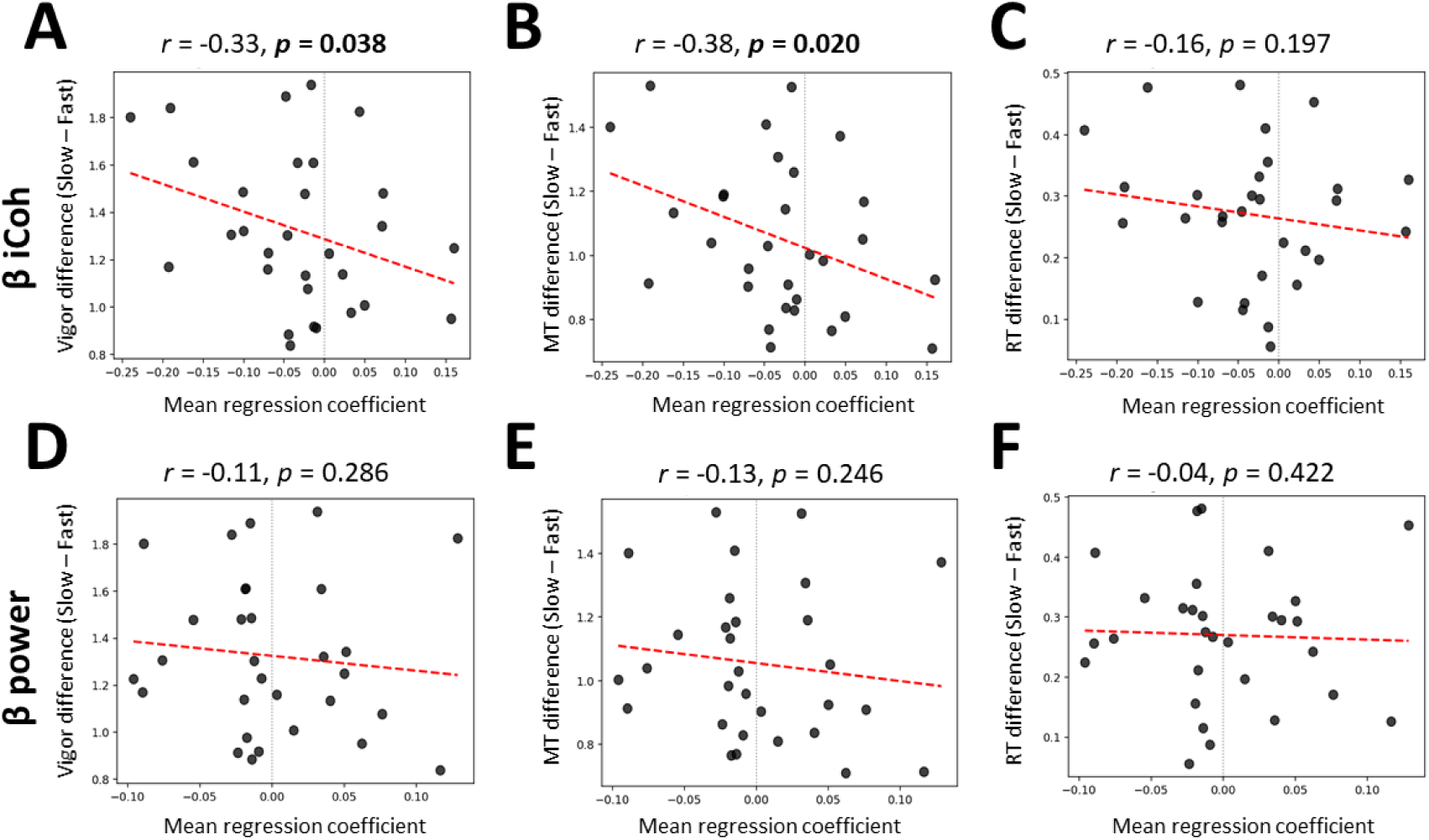
Between-subject correlation analysis of mean regression coefficients with vigor, MT and RT. Each panel illustrates the association found between mean regression coefficients based on β iCoh (first row) or β power (second row) and inter-individual differences in vigor (A, D), MT (B, E), and RT (C, F) between Fast and Slow conditions. Dark gray dots represent single-subject data and the regression line is depicted as a dotted red line. Pearson’s correlation coefficients and associated p-values are written on top of each of their corresponding plots.

## Discussion

The objective of this study was to determine if β power and iCoh, computed from high-density EEG data and studied across the whole scalp/brain in distinct sub-bands, can be used to accurately predict movement vigor. Participants executed movements at two instructed speeds (Fast, Slow), yielding significant differences in RT and MT, thereby manipulating vigor. Results showed that β power did not significantly differ between Fast and Slow conditions, whether computed over a pre-defined subset of left (contralateral to the moving hand) central electrodes or using subject-specific electrode selection. In contrast, β iCoh between left motor region and bilateral fronto-parietal regions was overall lower during Fast than Slow condition at the group level, both before and after go cue onset. Within-subject models incorporating subject-specific parcels selection revealed that changes in both β power and β iCoh could discriminate Fast and Slow trials better than chance in most participants (β power: 26/30, β iCoh: 28/30), but the direction of β power modulation greatly varied across subjects (∼50%/∼50% with decreased/increased β power in Fast in comparison to Slow conditions), while most participants (∼70%) showed lower β iCoh in Fast compared to Slow trials. Furthermore, β iCoh was not only predictive of vigor across trials, but also across subjects. Between-subject correlation analyses highlighted that regression coefficients based on β iCoh were negatively correlated with inter-individual Slow – Fast differences in vigor and MT, but not RT. No significant correlation was found between regression coefficients based on β power and inter-individual Slow – Fast differences in vigor, MT, and RT.

The absence of significant difference in left/contralateral motor β power when comparing fast and slow movements is consistent with results from previous studies [19,21–23]. A possible explanation of these findings is that movement-related β desynchronization is better linked to the integration of afferent sensory information than to efferent motor drive [43–45]. Yet, classification of Fast and Slow trials based on left motor β power achieved above-chance performance in most participants, suggesting some predictive value. However, the lateral/upper distribution of electrodes selected for these models, often excluding C3, indicates that peripheral artefacts may have influenced the results. This interpretation is further supported by the electrode distribution in models that included all electrodes, which was strongly biased toward peripheral scalp sites (although the most extreme prefrontal, temporal, and occipital electrodes were excluded). Since muscle activation is stronger and faster during rapid than slow movements, muscle artefacts could partly account for the above-chance performance of β power–based models. Indeed, muscle artefacts can introduce noise into the EEG signal at frequencies overlapping with the β band [46]. Importantly, restricting analyses to time windows preceding movement onset to minimize contamination by muscle artefacts did not improve model performance, nor did it reveal a consistent polarity of β power contrast between Fast and Slow conditions across participants. This suggests that movement vigor may not produce a consistent pattern of β power modulation across individuals. Such inter-individual variability has been documented in several studies [47,48], alongside evidence of the limited ability of motor β power to predict movement kinematics [7,21,45].

Classification of Fast and Slow trials based on β iCoh between the left motor cortex and bilateral fronto-parietal areas yielded above-chance performance in most participants with a majority exhibiting lower β iCoh in Fast than Slow conditions. The discriminative value of β iCoh extended beyond within-participant analyses: mean β iCoh regression coefficients were significantly negatively correlated with inter-individual vigor differences (Slow–Fast). Thus, participants with larger vigor difference showed more negative coefficients, reflecting greater reduction in β iCoh during Fast relative to Slow condition. No analogous relationship was observed for regression coefficients based on β power. Together, these findings indicate that β iCoh may constitute a more robust marker of movement vigor than β power, and that reducing β iCoh may increase vigor. Prior studies have reported scaling of contralateral motor gamma (50–100 Hz) power with movement speed [49] and a global facilitatory effect of gamma tACS on movement [17,50]. Gamma-band synchronization is thought to enable rapid formation of functional neural assemblies encoding motor commands [31,51]. A possible explanation, therefore, is that transient increases in motor cortical gamma activity are accompanied by decreased coherence (iCoh) at lower β frequencies with other cortical regions. Consistent with this idea, lower cortico-subcortical β coherence has been observed during movements compared with rest and associated with decreased RT [33,34]. To our knowledge, however, the present findings provide the first evidence of significant differences in cortico–cortical β iCoh according to movement speed. Furthermore, β iCoh was lower for fast than slow movements both before and after the go cue, suggesting that participants may have proactively downregulated β iCoh in anticipation of executing fast movements. This anticipatory control was likely facilitated by the block design, which allowed participants to know in advance the instructed speed.

The relatively stable β iCoh pattern identified here makes it a promising target for regulating movement vigor via non-invasive neurostimulation. Most available techniques, such as tACS or NF, are designed to modulate brain activity over several seconds and are therefore better suited to target sustained activity patterns. Downregulation of β iCoh during fast movements was also spatially broad, involving bilateral fronto-parietal regions. Rather than being a limitation, this spatial imprecision may represent an advantage for EEG-based neurostimulation, given the limited spatial resolution of EEG and the practical difficulty of precisely targeting individual brain areas. Nevertheless, more specific targeting could be achieved through source reconstruction combined with high-density EEG, as implemented in this study. Importantly, given the inter-individual variability in the fronto-parietal regions most predictive of movement speed, stimulation targets could be individualized by pre-identifying relevant regions with models based on calibration data. Similarly, the kernel used to project sensor-level signals into source space could be pre-computed on calibration data and then applied online to enable near–real time estimation of β iCoh at the source level. However, the feasibility and efficacy of such iCoh-based NF protocols for modulating movement vigor remain to be empirically tested.

Several limitations should be acknowledged. First, source reconstruction relied on a standard template rather than individual MRI data, meaning individual anatomical differences were not accounted for, which likely significantly reduced spatial accuracy at the individual scale. This may partly explain the inter-individual variability in the fronto-parietal regions whose β iCoh with left motor region best predicted movement speed. Still, this approach yielded above-chance performance in models predicting movement speed in most participants (28/30; 93.3%) and avoids the cost and limited accessibility of MRI. Second, the pre-movement period overlapped with a NF protocol targeting left motor β power, implemented for another study [37]. To minimize its influence, we pooled trials from opposite NF conditions (down- and up-regulation) along with passive/sham NF trials. The above-chance performance of most models trained with all of these trials, randomly shuffled across iterations, suggest that β iCoh predicted movement vigor independently of NF conditions. Moreover, as the last 1s before the go cue typically occurred late in the ∼8.5s NF window, participants often anticipated the go cue regardless of NF condition. Third, although β iCoh models achieved above-chance performance in most participants, classification accuracy was modest (mean AUC = 0.65). Additionally, while most participants exhibited negative regression coefficients, fewer than 30% were positive. Several factors may account for this variability, including residual single-trial noise, suboptimal parcel selection parameters, limited spatial resolution, or an intrinsically limited predictive value of β iCoh. Crucially, even within a deliberately simple modeling framework (logistic regression, minimal assumptions), β iCoh demonstrably outperformed local β power in predicting movement vigor, both within and across participants. This indicates that network-level β coupling carries vigor-relevant information not captured by local β power and motivates evaluation of alternative connectivity metrics and denser atlases.

## Conclusions

In conclusion, this study demonstrates that β iCoh between the left motor cortex and bilateral fronto-parietal regions may represent a more robust and consistent predictor of right-hand movement speed than left motor β power. Lower β iCoh predicted faster movements in both within-participant and across-participant analyses, indicating that network-level β coupling captures vigor-relevant information beyond local power. These findings identify β iCoh as a promising target for non-invasive neuromodulation to regulate movement vigor and as a feature for decoding intended movement speed in non-invasive BCIs, providing a more reliable alternative to β power-based approaches. Future work should assess the feasibility and efficacy of coherence-based stimulation and further explore the generalizability of connectivity-based metrics for predicting movement kinematics.

## Declarations

### Ethics approval and consent to participate

All participants gave their written informed consent before participation in the study. The study was approved by French ethical committee for the protection of individuals (CPP Nord-Ouest 1, ID RCB number nb. 2022-A00626-37) and conformed to the standards set by the latest version of the Declaration of Helsinki.

### Consent for publication

Not applicable.

### Availability of data and materials

The dataset analyzed in the current study is publicly available: https://doi.org/10.5281/zenodo.14638353

### Authors’ contributions

EP designed the experiment. CD and EP implemented the scripts for the experiment. CG and EP collected the data. APV developed scripts for source reconstruction and coherence computation. EP analyzed the data and wrote the first draft of the manuscript. CD, NG, CJK, and EP revised the manuscript.

