## Supplementary figures for "Cortical beta coherence provides a stronger non-invasive predictor of movement vigor than local beta power"

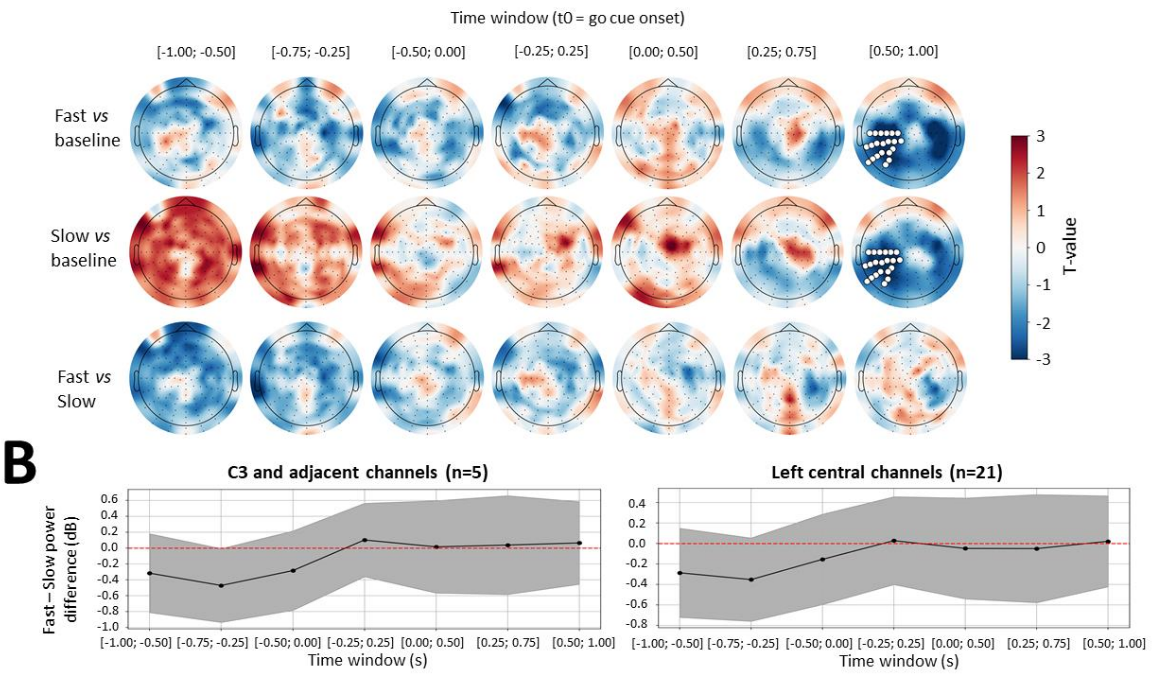


**Figure S1.** Analysis of mean difference in <u>low</u> β power between Fast and Slow conditions across the whole scalp and over left central electrodes. **A.** Topographical maps of t-values resulting from cluster-based permutation tests, which compared low β power in Fast condition and during baseline (top row), low β power in Slow condition and during baseline (middle row), and low β power in Fast and Slow conditions (bottom row). Mean difference in low β power was assessed in each 0.5s time window separately, that are indicated on top of each plot, in time relative to go cue. Electrodes that belonged to significant clusters ($p < 0.05$) are highlighted in white. **B.** Mean (dot) and 95% confidence intervals (shaded area) of the difference between Fast and Slow conditions in low β power, depicted per time window (x-axis), averaged over a cluster composed of C3 and adjacent electrodes (left) and over a larger cluster of left central electrodes centered on C3 (right). N indicates the number of electrodes included in each cluster.

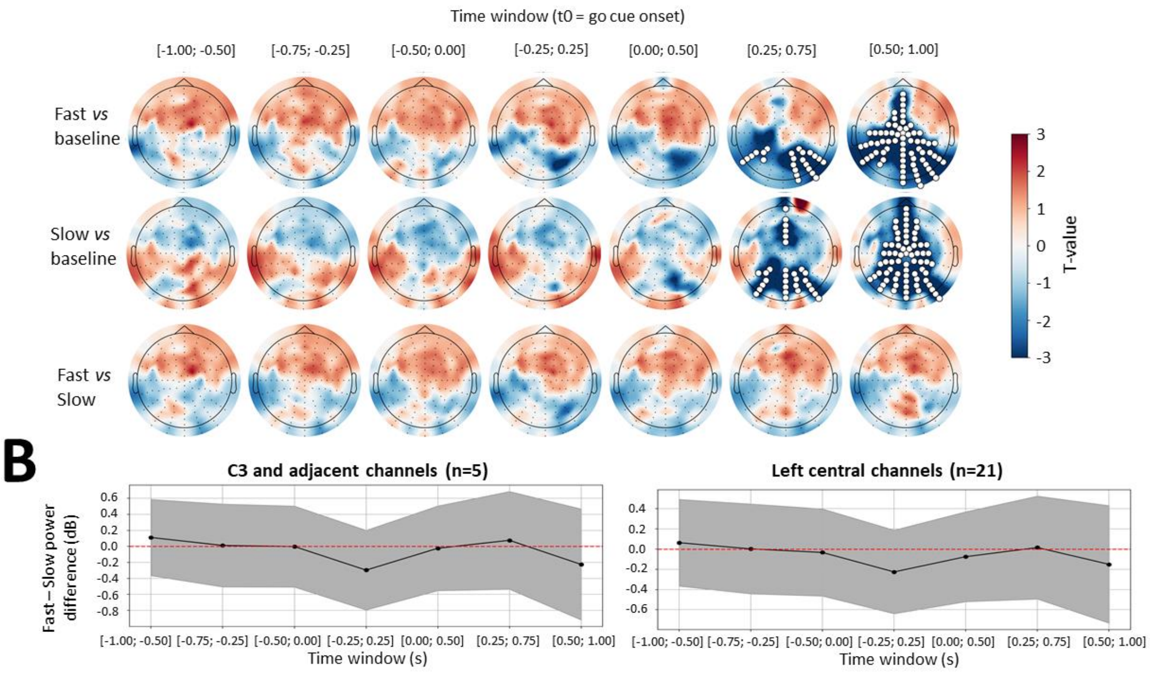


**Figure S2.** Analysis of mean difference in <u>high</u> β power between Fast and Slow conditions across the whole scalp and over left central electrodes. **A.** Topographical maps of t-values resulting from cluster-based permutation tests, which compared high β power in Fast condition and during baseline (top row), high β power in Slow condition and during baseline (middle row), and high β power in Fast and Slow conditions (bottom row). Mean difference in high β power was assessed in each 0.5s time window separately, that are indicated on top of each plot, in time relative to go cue. Electrodes that belonged to significant clusters ($p < 0.05$) are highlighted in white. **B.** Mean (dot) and 95% confidence intervals (shaded area) of the difference between Fast and Slow conditions in high β power, depicted per time window (x-axis), averaged over a cluster composed of C3 and adjacent electrodes (left) and over a larger cluster of left central electrodes centered on C3 (right). N indicates the number of electrodes included in each cluster.

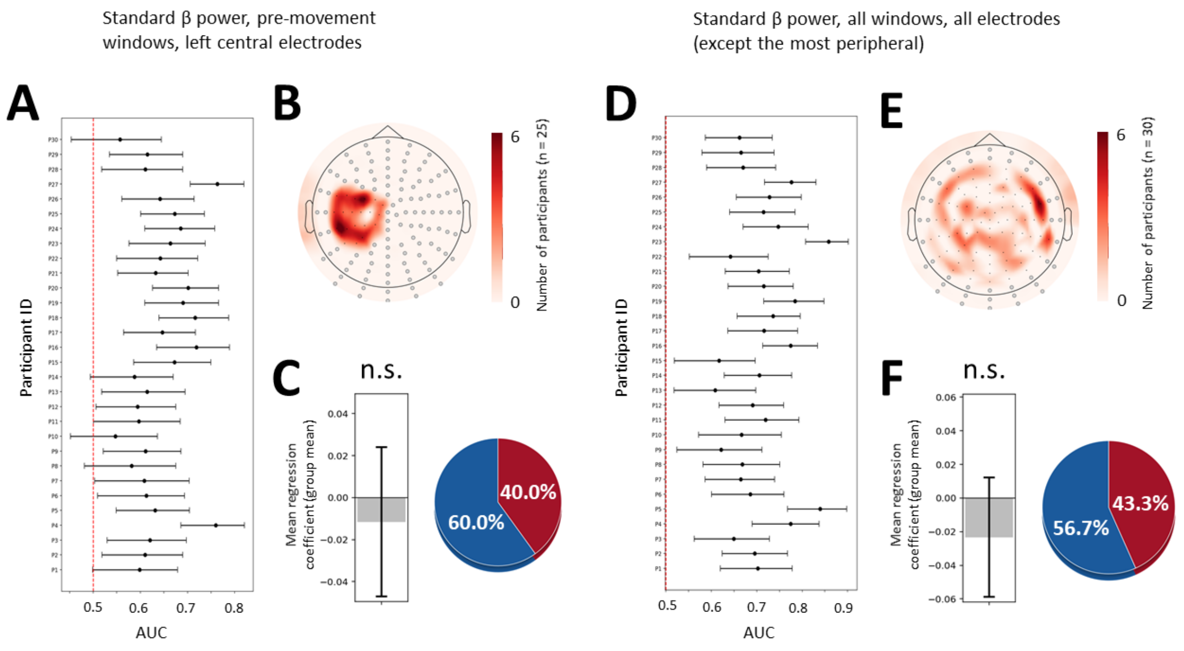


**Figure S3.** Analysis of mean difference in β power between Fast and Slow conditions with subject-specific selection of left central electrodes in pre-movement time windows (A, B, C) and all electrodes (except peripheral) in all time windows (D, E, F). **A, D.** 95% confidence intervals around the mean AUC represented as error bars for each participant (y-axis) resulting from logistic regression models of speed instruction (Fast vs Slow) based on β power. The red dotted line indicates chance level (AUC = 0.5). **B, E.** Scalp topography of electrodes according to their selection frequency. Red areas indicate electrodes most often selected across participants. Excluded electrodes appear as filled gray circles. **C, F.** Left, mean regression coefficient (averaged across subjects) represented as a gray bar, with the error bar illustrating 95% confidence interval. N.s. = not significant ($p > 0.05$). Right, pie chart illustrating the percentage of participants showing positive and negative regression coefficients in red and blue, respectively.

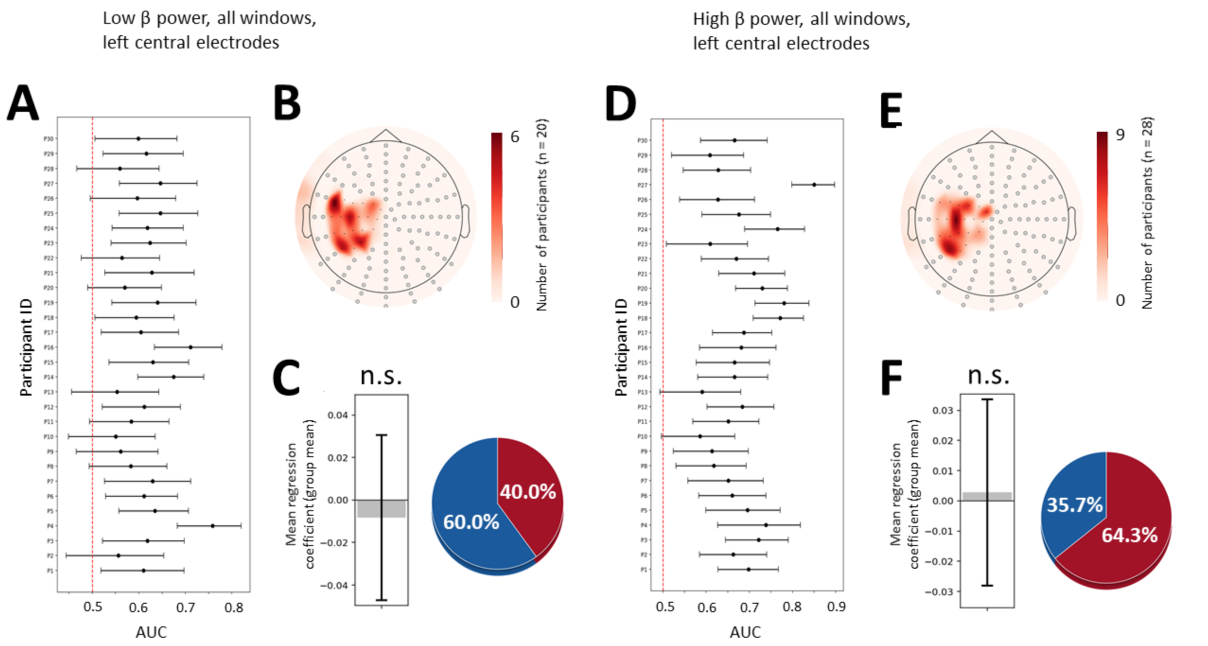


**Figure S4.** Analysis of mean difference in low β power (A, B, C) and high β power (D, E, F) between Fast and Slow conditions with subject-specific selection of left central electrodes. **A, D.** 95% confidence intervals around the mean AUC represented as error bars for each participant (y-axis) resulting from logistic regression models of speed instruction (Fast vs Slow) based on β power. The red dotted line indicates chance level (AUC = 0.5). **B, E.** Scalp topography of electrodes according to their selection frequency. Red areas indicate electrodes most often selected across participants. Excluded electrodes appear as filled gray circles. **C, F.** Left, mean regression coefficient (averaged across subjects) represented as a gray bar, with the error bar illustrating 95% confidence interval. N.s. = not significant ($p > 0.05$). Right, pie chart illustrating the percentage of participants showing positive and negative regression coefficients in red and blue, respectively.

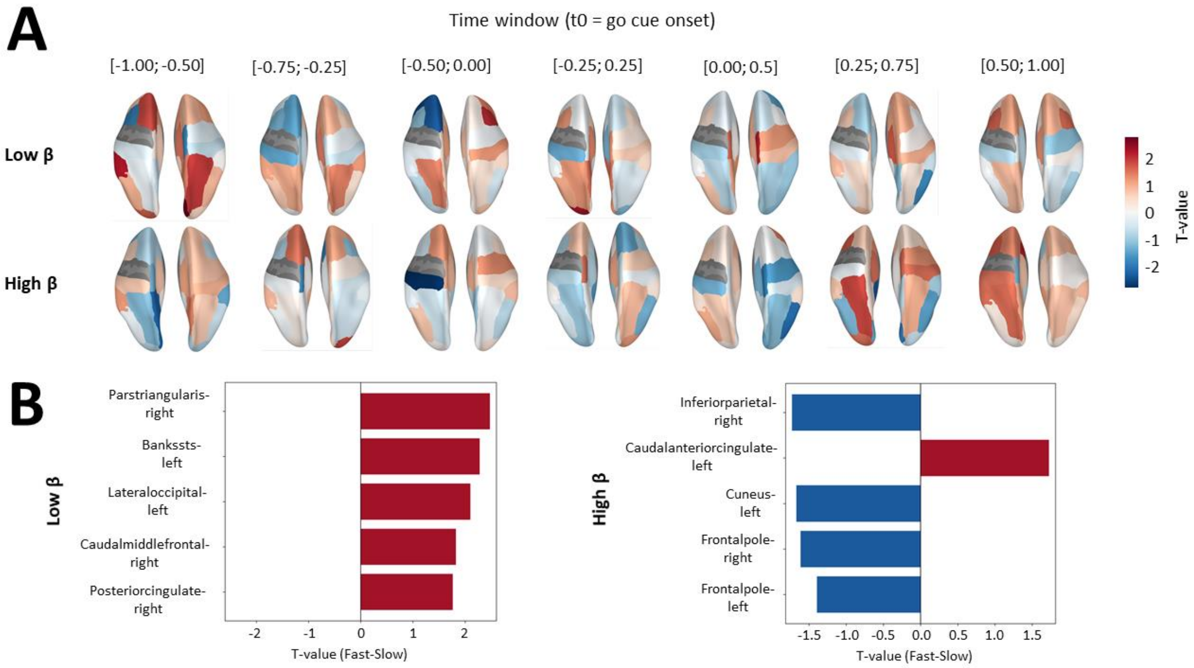


**Figure S5.** Analysis of mean difference in low and high β iCoh between Fast and Slow conditions across parcels. **A.** Topographical maps of t-values resulting from the comparison of low β (top row) and high β (bottome row) iCoh in Fast and Slow conditions. Mean difference in β power was assessed in each 0.5s time window separately, that are indicated on top of each plot, in time relative to go cue. Hot and cold colors respectively indicate higher (positive t-value) and lower (negative t-value) β iCoh in Fast than Slow conditions in average. **B.** Top five parcels with the highest absolute t-value averaged across time windows for low β (left panel) and high β (right panel) iCoh. Red and blue bars respectively indicate parcels associated with a positive and negative t-value.

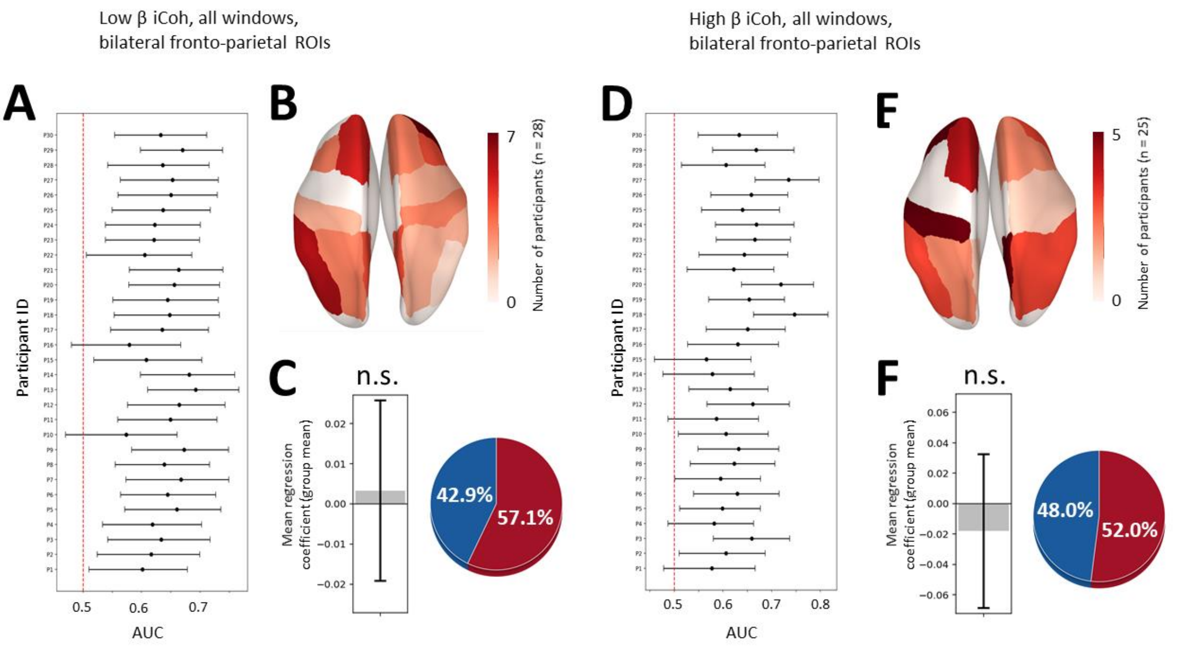


**Figure S6.** Analysis of mean difference in low β (A, B, C) and high β iCoh (D, E, F) between Fast and Slow conditions with subject-specific selection of bilateral fronto-parietal parcels. **A, D.** 95% confidence intervals around the mean AUC represented as error bars for each participant (y-axis) resulting from logistic regression models of speed instruction (Fast vs Slow) based on low β (A) and high β (D) iCoh. The red dotted line indicates chance level (AUC = 0.5). **B, E.** Brain topography of fronto-parietal parcels according to their selection frequency. Red areas indicate parcels most often selected across participants. **C, F.** Left, mean regression coefficient (averaged across subjects) represented as a gray bar, with the error bar illustrating 95% confidence interval. N.s. = not significant ($p > 0.05$). Right, pie chart illustrating the percentage of participants showing positive and negative regression coefficients in red and blue, respectively.

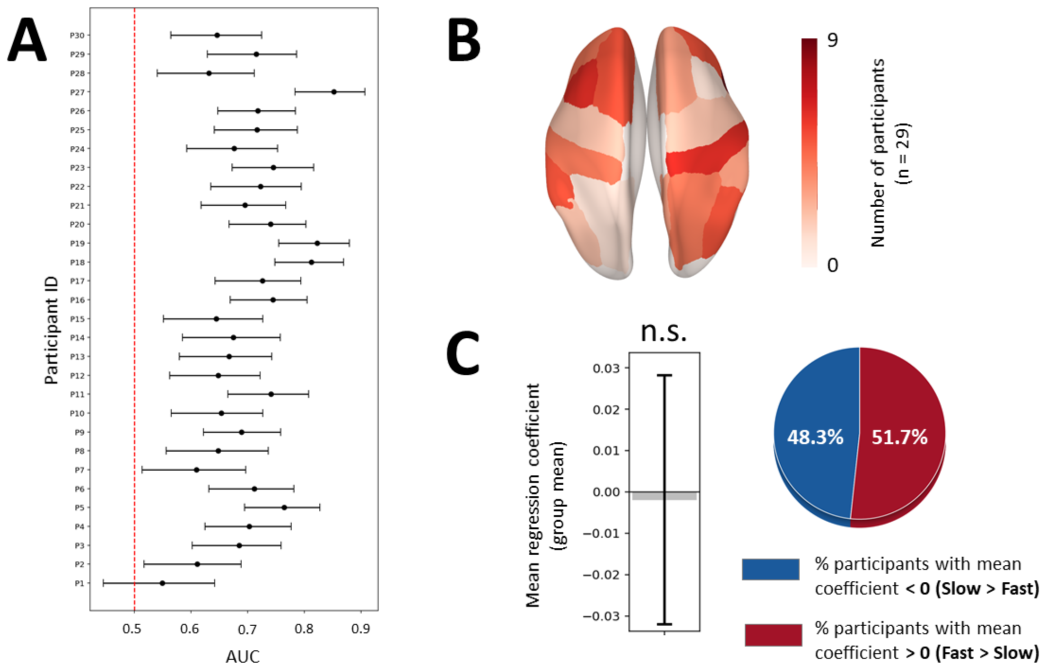


**Figure S7.** Analysis of mean difference in source-level β power between Fast and Slow conditions with subject-specific selection of bilateral fronto-parietal parcels. **A, D.** 95% confidence intervals around the mean AUC represented as error bars for each participant (y-axis) resulting from logistic regression models of speed instruction (Fast vs Slow) based on β iCoh. The red dotted line indicates chance level (AUC = 0.5). **B, E.** Brain topography of fronto-parietal parcels according to their selection frequency. Red areas indicate parcels most often selected across participants. **C, F.** Left, mean regression coefficient (averaged across subjects) represented as a gray bar, with the error bar illustrating 95% confidence interval. N.s. = not significant ($p > 0.05$). Right, pie chart illustrating the percentage of participants showing positive and negative regression coefficients in red and blue, respectively.
